# Genetic diversity of *Lactobacillus delbrueckii* in home-made *Dahi* from different regions of India as revealed by genotyping-by-sequencing

**DOI:** 10.64898/2026.01.09.698673

**Authors:** Pragya Tripathi, Vivek Kumar Singh, Mahesh Tiwari, Shashi Bhushan Tripathi

**Affiliations:** Independent Researcher; Department of Biotechnology, TERI School of Advanced Studies, 10, Institutional Area, Vasant Kunj, New Delhi, India 110070; Department of Agriculture, National Post-graduate College, Barhalganj, Gorakhpur, India 273402

**Keywords:** Probiotics, Lactic acid bacteria, Genetic diversity, food safety

## Abstract

Microbial diversity of bacterial species in home-made curd from different states of India was analyzed using genotyping-by-sequencing. The most prevalent species were *Lactobacillus delbrueckii* subsp. *bulgaricus, Lactobacillus helveticus, Streptococcus thermophilus* and *Lactobacillus kefiranofaciens*. Pathogenic species such as *Salmonella enterica* serovar *Typhimurium* and *Shigella dysenteriae* were found in over 60% of samples. High intraspecific variation was observed in *Acinetobacter baumannii, Bacillus cereus, Enterococcus faecium, Escherichia coli, Lactococcus lactis* subsp. *cremoris, Lactococcus piscium, Salmonella enterica* serovar *Typhimurium* and *Shigella dysenteriae*. Intra-specific genetic relatedness in *Lactobacillus delbrueckii* subsp. *bulgaricus* in curd samples largely correlated with their respective sites of collection.

**Research Highlights:**

- Metagenomic DNA from home-made Dahi (curd) samples from various states in India were used for GBS-SNPs
- GBS-SNPs were able to resolve within-species variability across the samples.
- Majority of the samples contained both probiotic as well as potential pathogenic bacteria.
- Several of curd samples contained multiple strains.

## 1. Introduction

Fermented foods constitute an important part of the human diet worldwide. Curd (hereafter mentioned as *dahi*) is a traditional Indian fermented dairy product. *Dahi* finds its mention in the oldest Hindu scripts and remains an indispensable part of all cultural and religious rituals. The consumption of industrially produced *dahi* has been consistently increasing in India, however, home-made *dahi* has a major share, especially in rural India where *dahi* making is a daily household activity. Traditionally, *dahi* is prepared using raw or boiled milk followed by inoculation through back-slopping and incubation for 5-6 hours or overnight as per requirement (Joishy *et al*. 2019). Under household settings, the incubation temperature, in general, is not controlled and is left at ambient temperature. This natural fermentation process leads to wide variations in microbial composition and sensory quality of the product unlike those of industrial products where pure cultures and controlled incubation temperatures are used to maintain a uniform product quality. Consequently, the probiotic and sensory properties of such home-made *dahi* may differ with geographical locations due to environmental variations which affect the overall bacterial microflora (Zhao *et al*. 2021). In addition to the species composition, home-made *dahi* from different geographical localities may also harbour genetically distinct strains of *Lactobacillus* and other milk-fermenting bacteria which may impart a distinct flavour and taste to the product. On the other hand, these home-made products are also prone to undesirable contamination with hazardous microbes (Powell *et al*. 2011).

Most common probiotic bacteria in curd are *Lactobacillus acidophilus, Lactobacillus johnsonii, Lacticaseibacillus casei, Streptococcus thermophilus, Lactobacillus helveticus, Lactococcus lactis, Lactococcus cremoris, Lactobacillus delbrueckii, Lactobacillus rhamnosus, Limosilactobacillus fermentum, Enterococcus faecium, Leuconostoc mesenteroides, Lactobacillus plantarum, Lactobacillus kefiranofaciens* and *Lacticaseibacillus paracasei* (Farahmand *et al*. 2021; Islam *et al*. 2021; Jayashree *et al*. 2013). However, improper handling or non-sterile conditions during fermentation can lead to the growth of other bacteria also. These may sometimes include pathogens such as *Streptococcus infantarius, Clostridium spp*., *Escherichia coli, Salmonella typhimurium, Enterococcus faecalis, Listeria monocytogenes* and *Staphylococcus aureus* (Negi *et al*. 2018; Sessou *et al*. 2023).

Characterisation of bacterial communities in fermented foods is traditionally done using culture-based methods. However, this approach is considered ineffective for several bacteria which are “unculturable” (Didelot *et al*. 2012). In the past few decades, advances in metagenomics have revolutionized the study of microbial communities (Jayashree *et al*. 2013; Kopera *et al*. 2024). Amplicon sequencing of conserved genomic regions of 16S rRNA in bacteria, using next generation sequencing, is the most widely used approach to characterize microbial communities due to its lower cost. Alternatively, whole genome shotgun sequencing is used which has higher cost but allows a more precise taxonomic and functional classification of sequences including strain-level differentiation (Jovel *et al*. 2016). Whole genome shotgun sequencing also allows for differentiation of bacterial strains which is not possible with 16S rRNA sequencing.

Genotyping by sequencing (GBS-SNP) is a high throughput genotyping technique based on next generation sequencing of reduced representation genomic libraries (Elshire *et al*. 2011). The technique randomly samples nucleotide-level variations from a genome-wide context like shotgun sequencing, although its coverage and cost are much lesser. We propose that the GBS-SNPs can be used to detect the present/absence of any target set of species, without isolation of individual species from each sample, by mapping the sequence reads to those bacterial genome assemblies. More importantly, GBS-SNPs can also provide strain-level resolution like shotgun sequencing at a significantly lower cost.

In this study, we used GBS-SNPs to investigate a pre-determined set of bacterial species in home-made dahi samples from geographically distant locations of India for their occurrence. Further, we characterized the intra-specific genetic diversity of one of the most frequent species, *Lactobacillus delbrueckii* subsp. *bulgaricus* across different states of India.

## 2. Materials & Methods

### 2.1. Collection of dahi samples

Thirty-three *dahi* samples were collected from geographically distant locations in India (Supplementary table S1). All samples except sample L from Delhi were collected from local households where *dahi* making was a daily activity. The samples were brought to laboratory under chilling condition and stored at -20 °C till isolation of total DNA.

### 2.2. Isolation of DNA

Genomic DNA was isolated from *dahi* samples using a modified CTAB based method (Rana *et al*. 2023). In brief, each sample was diluted 1:1 with TNE buffer and was centrifuged at 10000 RPM for 10 minutes. The pellet was resuspended in TNE buffer and recovered through centrifugation. The pellet was resuspended in 300 ml of TNE buffer and an equal volume of 2x CTAB buffer was added and incubated at 65 °C for 30 minutes. The lysate was extracted with chloroform:isoamyl alcohol (C:I), treated with RNAse A (50 μg/ml) for 30 minutes at room temperature, and reextracted with C:I. High molecular weight DNA was precipitated with isopropanol and recovered by centrifugation. The DNA pellet was washed with 70% ethanol, air-dried, and finally dissolved in 50 μl of TE buffer.

### 2.3. GBS-SNP Library preparation and sequencing

GBS-SNP libraries were prepared using 100 ng of DNA from each sample following Elshire *et al*., (2011) with only modification that the restriction enzyme, PstI, was used instead of ApeK1. Briefly, individual DNA samples were digested with PstI, following which the barcoded PstI adapters were ligated. The individual ligation reactions were pooled, and the pooled library was purified using Monarch® PCR & DNA Cleanup Kit (New England Biolabs, MA, USA). The purified library was then amplified in 50-µl reaction volumes using 5 µl of the purified library as template DNA with 5x PCR Mix (New England Biolabs, MA, USA) with following primers: 5′-AATGATACGGCGACCACCGAGATCTACACTCTTTCCCTACACGACGCTCTTCCGATCT-3′ and 5′-CAAGCAGAAGACGGCATACGAGATCGGTCTCGGCATTCCTGCTGAACCGCTCTTCCG ATCT-3′

Thermal cycling parameters consisted of 72 °C for 5 min, 95 °C for 30 s followed by 18 cycles of 95 °C for 30 s, 65 °C for 10 s, and 72 °C for 30 s, with a final extension step at 72 °C for 5 min. The purified GBS-SNP library was sequenced at 150 bp x 2 on Novaseq X using a commercial sequencing facility (Macrogen, South Korea).

### 2.4. SNP calling and data analysis

In this study, we selectively targeted twenty-one bacterial species which included probiotic, pathogenic and spoilage-related species reported to be found in *dahi* (Table 1). Reference genome assemblies of these 21 bacteria were used for mapping of the GBS-SNP reads and SNP calling. Eighteen of the reference genome assemblies were chromosome-level assemblies and remaining three genome assemblies were available in the form of multiple scaffolds (Supplementary table S2). Processing of FASTQ files containing raw sequence reads was done using the Linux-based GBS analysis pipeline as implemented in the program TASSEL v5.0 (Glaubitz *et al*. 2014). Unique sequence tags were aligned to bacterial reference genome assemblies using Bowtie2 available as a web-based tool at Galaxy platform (https://usegalaxy.org/). GBS-SNPs were called in the variant calling format (VCF).

**Table 1.**
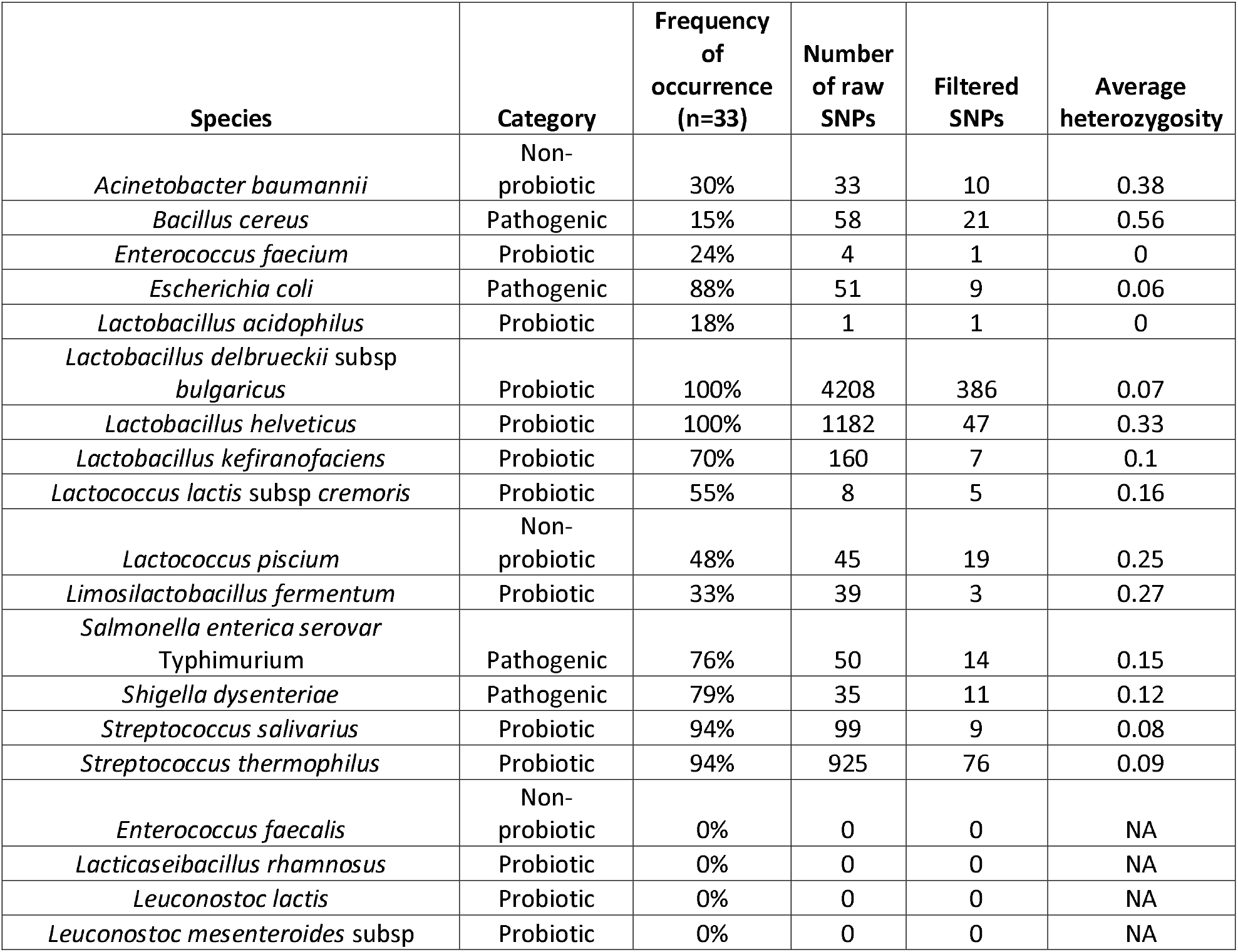
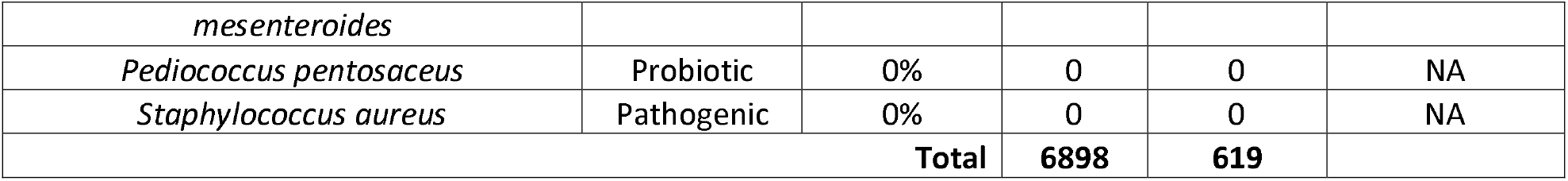
Number of mapped GBS-SNPs and observed heterozygosity in bacterial species in *dahi* samples.

The VCF file containing GBS-SNPs was processed using GUI version of TASSEL v5.0 (Glaubitz *et al*. 2014). For each sample, the presence or absence of bacterial species was inferred from the GBS-SNPs mapping to the respective reference genomes. Complete absence of GBS-SNP from a particular reference genome in a sample was considered to imply absence of the species in that sample. Consequently, a binary matrix (1 for presence, 0 for absence) was generated for each sample. Genetic relatedness of *Lactobacillus delbrueckii* subsp. *bulgaricus* was analysed from high quality filtered GBS-SNP data by using a single SNP per read and including SNPs only with less than 50% of missing data.

## 3. Results and discussion

### 3.1. Probiotic and pathogenic bacteria in dahi samples

DNA from a total of 33 *dahi* samples was used for GBS-SNP genotyping. The number of GBS reads per sample varied from 50409 (in P3) to 30909537 (in U2) with an average of 9.1 million reads per sample (see supplementary material). Thirty of the samples had greater than 1 million reads per sample. A total of 6898 GBS-SNPs were initially obtained from these reads. Subsequently, only one GBS-SNP per read was considered which resulted in 619 high quality GBS-SNPs (Table 1). These GBS-SNPs were found to be mapping to reference genomes of 15 out of 21 bacterial species used for SNP calling. Such mapping of GBS reads to a particular reference genome assembly was assumed to be indicative of presence of the species in that *dahi* sample. No GBS-SNPs were found to be mapping on remaining 6 reference genomes in any sample, which implied that those species were absent in the samples. These species were *Enterococcus faecalis, Lacticaseibacillus rhamnosus, Leuconostoc lactis, Leuconostoc mesenteroides* subsp. *mesenteroides, Pediococcus pentosaceus* and *Staphylococcus aureus*. It may be possible that these species were present at levels undetectable using GBS-SNP genotyping.

The frequency at which the 15 bacterial species occurred in *dahi* samples varied from 15% to 100% (Figure 1 and Supplementary table S3). Four of the most frequently detected species belonged to probiotic category. These were, *Lactobacillus helveticus* (100%), *Lactobacillus delbrueckii* subsp. *bulgaricus* (100%), *Streptococcus salivarius* (94%), *Streptococcus thermophilus* (94%) and *Escherichia coli* (88%). Our results resonate very well with those from earlier studies where *Lactobacillus delbrueckii* subsp. *bulgaricus*, and *Streptococcus thermophilus* were the major bacteria in sour yoghurt (Islam *et al*. 2021). The least frequently observed species were *Bacillus cereus* (15%) and *Lactobacillus acidophilus* (18%) which were detected in 15% and 18% of all *dahi* samples only.

**Fig. 1.**
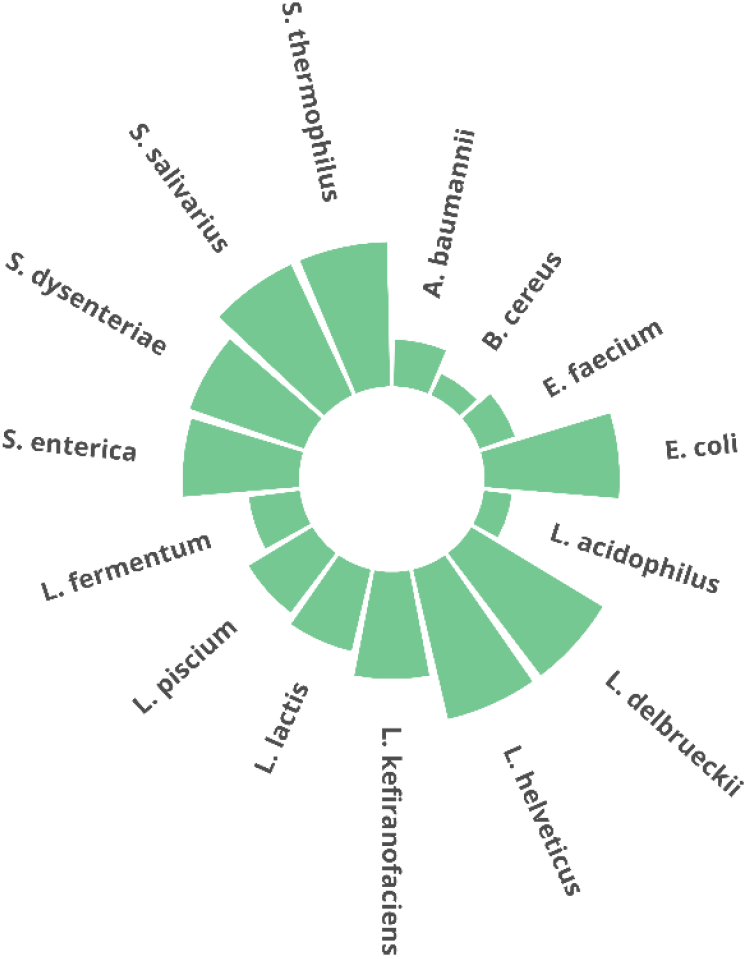
Frequency of occurrence of fifteen bacterial species in 33 *dahi* samples as revealed by GBS-SNPs

The bacteria detected in *dahi* samples also included four pathogenic species. Three of these, namely, *Escherichia coli, Shigella dysenteriae* and *Salmonella enterica* serovar Typhimurium were detected in 88%, 79% and 76% of the samples, respectively. The results indicate that these bacteria are very frequent in home-made *dahi*. There was no correlation between the occurrence of these bacteria and geographical locations of samples. High prevalence of *E. coli* contamination in samples of raw milk (47%) have been reported from South India (Joseph and Kalyanikutty 2022). However, there was no indication of any negative effect of these species on the household members. It is possible that the pathogenic effect of these species in *dahi* is suppressed by more prevalent probiotic species. Growth inhibition of *Shigella dysenteriae* and *Salmonella enterica* serovar Typhimurium by yoghurt has been reported (Alm 1983). Yoghurt samples inoculated with staphylococci up to 10^5^ cells per gram, were shown to be free of viable staphylococci by the 2^nd^ to 4^th^ day of storage which may be due to dominance of other bacteria present in the starter (Pazakova *et al*. 1997).

All *dahi* samples contained multiple bacterial species as revealed by the GBS-SNPs. Sixteen of the samples contained 10 or more species (Supplementary table S3). Overall, the number of species among the samples varied from 4 (in VMS5 from Varanasi) to 13 (in TN1 from Chennai) with an average of 9.2 species per sample. The number of species in the samples did not correlate with their geographical origin. This may be because the bacterial composition of a *dahi* sample is likely to be influenced by multiple factors such as the bacterial composition of the starter, handling of raw materials, and growth conditions during incubation of the product. These factors may vary greatly among households even within a locality.

### 3.2. Genetic diversity in Lactobacillus delbrueckii subsp bulgaricus

Out of 619 GBS-SNPs obtained after filtering, 386 GBS-SNPs mapped to *Lactobacillus delbrueckii* subsp *bulgaricus* reference genome. GBS-SNP data from three of the *dahi* samples, namely, K1, K4 and O1, was excluded from further analysis as these had high amount of missing GBS-SNP data. The SNP data was used to calculate pair-wise genetic distances among the samples (supplementary table S4). The maximum genetic distance among the samples was 0.385 (between VMS3 and V2 from Varanasi) with an average genetic distance of 0.132. The neighbour-joining dendrogram showed three major clusters (Figure 2). Cluster 1 contained five out of six samples from Delhi and all the three samples from Tamil Nadu forming a relatively tight cluster. Six out of nine samples from Uttar Pradesh grouped together in the cluster 2 which also contained 3 out of 7 samples from Rajasthan. Cluster 3 was the most diverse and included both samples from Uttarakhand, both the samples from Maharashtra, 4 out of 7 samples from Rajasthan, 3 samples from Uttar Pradesh and one sample each from Delhi and Odisha. It may be noted that the sample L from Delhi belonged to an industrial brand and was not home-made. Thus, clustering of samples largely correlated with their locality of collection with some exceptions. It is possible that distinct strains of this species are prevalent in different localities.

**Fig. 2.**
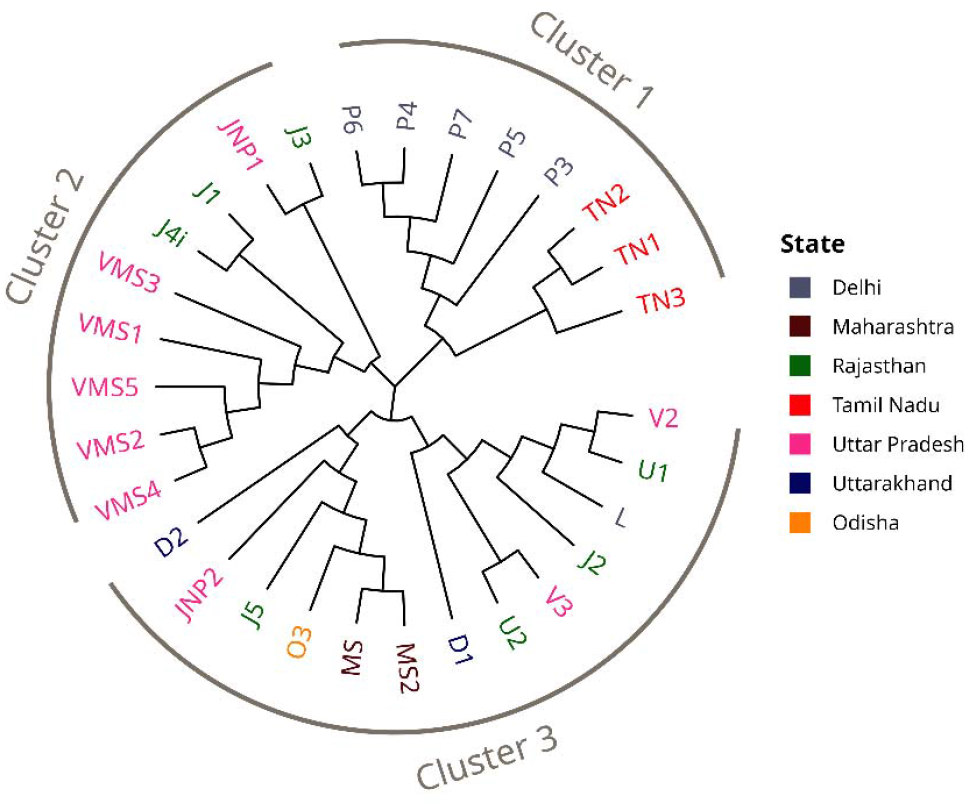
Neighbour-joining tree showing genetic relatedness of *Lactobacillus delbrueckii* subsp. *bulgaricus* in *dahi* samples

It was interesting to note that several of samples contained multiple strains of a particular species which was indicated by the presence of heterozygous GBS-SNP sites. Considering the haploid nature of bacterial genomes, all the sites are expected to be homozygous, and a heterozygous SNP call is possible only if two different strains are present in the sample. Heterozygous sites were observed for all species except *Enterococcus faecium* and *Lactobacillus acidophilus*. The proportion of heterozygous sites in the data set varied from 6% to 56% with an overall heterozygous proportion of 17% (Table 1). It was interesting to note that a higher level of heterozygosity was observed in *Acinetobacter baumannii, Bacillus cereus, Enterococcus faecium, Escherichia coli, Lactococcus lactis* subsp. *cremoris, Lactococcus piscium, Salmonella enterica* serovar *Typhimurium* and *Shigella dysenteriae*. On the other hand, in *Lactobacillus delbrueckii* subsp. *bulgaricus, Streptococcus salivarius* and *Streptococcus thermophilus*, which were the most prevalent probiotic species in this study, heterozygosity levels remained relatively low at about 7-8%. Low heterozygosity in these may indicate a relatively lower genetic variability within the species at a local level. Nevertheless, our data shows that home-made *dahi* samples harbour microbial diversity in the form of multiple species as well as in the form of multiple strains of a species.

GBS-SNPs have been widely used for genetic diversity studies in various plant and animal species. However, it has been used less frequently to study diversity in microbial communities. The metagenomic approach using 16S rRNA are the best suitable for studying the genus or species level microbial composition of samples. However, due to high level of sequence conservation and low genome coverage, they cannot distinguish strains. The GBS-SNPs due their higher genome coverage through hundreds of randomly spread SNPs, are best suitable for strain-level differentiation and investigation of genetic relatedness. This study also shows that GBS-SNPs can be used on environmental samples containing multiple species without a need for isolation of the strains. The list of target species can be changed as per requirement by simply providing the target reference genome(s) during the read alignment and SNP calling.

## 4. Conclusion

This study shows that GBS-SNPs can provide useful species and strain-level resolution in samples containing multiple species. The study showed that home-made *dahi* commonly harbours multiple bacterial species including those considered as pathogenic. GBS-SNPs could also provide strain-level resolution and information on relatedness of strains in case of *Lactobacillus delbrueckii* subsp. *bulgaricus*.

## Abbreviations

GBS: Genotyping by sequencing
SNP: Single nucleotide polymorphism
VCF: Variant call format
GUI: Graphical user interface
CTAB: Cetyltrimethylammonium Bromide;

## Acknowledgements

The authors thank the student volunteers for their help in collection of samples from diverse locations. We acknowledge TERI School of Advanced Studies for providing support for research infrastructure.

## Authors’ contributions

Pragya Tripathi – Methodology, Bioinformatic analysis, Map visualization, Writing – original draft, Writing – review & editing

Vivek Kumar Singh – Methodology, Writing – review & editing

Mahesh Tiwari-Methodology, Supervision

Shashi Bhushan Tripathi – Resources, Supervision, Conceptualization, Writing – original draft, Writing – review & editing

## References

Alm, L., 1983. Survival rate of Salmonella and Shigella in fermented milk products with and without added human gastric juice: an in vitro study. Progress in Food & Nutrition Science. 7, 19–28,

Didelot, X., Bowden, R., Wilson, D.J., Peto, T.E.A., Crook, D.W., 2012. Transforming clinical microbiology with bacterial genome sequencing. Nature Rev. Genet. 13, 601–612, 10.1038/nrg3226

Elshire, R.J., Glaubitz, J.C., Sun, Q., Poland, J.A., Kawamoto, K., Buckler, E.S., Mitchell, S.E., 2011. A robust, simple genotyping-by-sequencing (GBS) approach for high diversity species. PloS One. 6, e19379,

Farahmand, N., Ouoba, L.I.I., Naghizadeh Raeisi, S., Sutherland, J., Ghoddusi, H.B., 2021. Probiotic Lactobacilli in fermented dairy products: Selective detection, enumeration and identification scheme. Microorganisms. 9, 10.3390/microorganisms9081600

Glaubitz, J.C., Casstevens, T.M., Lu, F., Harriman, J., Elshire, R.J., Sun, Q., Buckler, E.S., 2014. TASSEL-GBS: a high capacity genotyping by sequencing analysis pipeline. PloS One. 9, e90346,

Islam, S.M.R., Tanzina, A.Y., Foysal, M.J., Hoque, M.N., Rumi, M.H., Siddiki, A., Tay, A.C., Hossain, M.J., Bakar, M.A., Mostafa, M., Mannan, A., 2021. Insights into the nutritional properties and microbiome diversity in sweet and sour yogurt manufactured in Bangladesh. Scientific Reports. 11, 22667, 10.1038/s41598-021-01852-9

Jayashree, S., Pushpanathan, M., Rajendhran, J., Gunasekaran, P., 2013. Microbial diversity and phylogeny analysis of buttermilk, a fermented milk product, employing 16S rRNA-based pyrosequencing. Food Biotechnol. 27, 213–221, 10.1080/08905436.2013.811084

Joishy, T.K., Dehingia, M., Khan, M.R., 2019. Bacterial diversity and metabolite profiles of curd prepared by natural fermentation of raw milk and back sloping of boiled milk. World J. Microbiol. Biotechnol. 35, 102, 10.1007/s11274-019-2677-y

Joseph, J., Kalyanikutty, S., 2022. Occurrence of multiple drug-resistant Shiga toxigenic Escherichia coli in raw milk samples collected from retail outlets in South India. J. Food Sci.Technol. 59, 2150–2159, 10.1007/s13197-021-05226-x

Jovel, J., Patterson, J., Wang, W., Hotte, N., O’Keefe, S., Mitchel, T., Perry, T., Kao, D., Mason, A.L., Madsen, K.L., Wong, G.K., 2016. Characterization of the gut microbiome using 16S or shotgun metagenomics. Front. Microbiol. 7, 459, 10.3389/fmicb.2016.00459

Kopera, K., Gromowski, T., Wydmański, W., Skonieczna-Żydecka, K., Muszyńska, A., Zielińska, K., Wierzbicka-Woś, A., Kaczmarczyk, M., Kadaj-Lipka, R., Cembrowska-Lech, D., Januszkiewicz, K., Kotfis, K., Witkiewicz, W., Nalewajska, M., Feret, W., Marlicz, W., Łoniewski, I., Łabaj, P.P., Rydzewska, G., Kosciolek, T., 2024. Gut microbiome dynamics and predictive value in hospitalized COVID-19 patients: a comparative analysis of shallow and deep shotgun sequencing. Front. Microbiol. 15, 1342749, 10.3389/fmicb.2024.1342749

Negi, Y.K., Pandey, C., Saxena, N., Sharma, S., Garg, F.C., Garg, S.K., 2018. Isolation of antibacterial protein from Lactobacillus spp. and preparation of probiotic curd. J. Food Sci.Technol. 55, 2011–2020, 10.1007/s13197-018-3115-0

Pazakova, J., Turek, P., Laciakova, A., 1997. The survival of Staphylococcus aureus during the fermentation and storage of yoghurt. J. Appl. Microbiol. 82, 659–662, 10.1111/j.1365-2672.1997.tb03599.x

Powell, I.B., Broome, M.C., Limsowtin, G.K.Y. In: Fuquay JW, editor Encyclopedia of Dairy Sciences (Second Edition). San Diego: Academic Press, pp. 552–558, 2011.

Rana, A., Malik, A.A., Tripathi, S.B., Kumar, A., 2023. Novel SNP based analysis of genetic diversity in Polygonatum verticillatum Linn. across Indian Himalayas. 3 Biotech. 13, 242, 10.1007/s13205-023-03654-4

Sessou, P., Keisam, S., Gagara, M., Komagbe, G., Farougou, S., Mahillon, J., Jeyaram, K., 2023. Comparative analyses of the bacterial communities present in the spontaneously fermented milk products of Northeast India and West Africa. Front. Microbiol. 14, 10.3389/fmicb.2023.1166518

Zhao, Z., Ning, C., Chen, L., Zhao, Y., Yang, G., Wang, C., Chen, N., Zhang, Z., Li, S., 2021. Impacts of manufacture processes and geographical regions on the microbial profile of traditional Chinese cheeses. Food Research International. 148, 110600, 10.1016/j.foodres.2021.110600

