## Supplementary material for "Genetic diversity of *Lactobacillus delbrueckii* in home-made *Dahi* from different regions of India as revealed by genotyping-by-sequencing"

**Table S1.** Details of *dahi* samples used in the study

| **SampleID** | **State** | **City** |
| --- | --- | --- |
| L | Delhi | New Delhi |
| P3 | Delhi | New Delhi |
| P4 | Delhi | New Delhi |
| P5 | Delhi | New Delhi |
| P6 | Delhi | New Delhi |
| P7 | Delhi | New Delhi |
| K1 | Kerala | Kollam |
| K4 | Kerala | Kollam |
| MS | Maharashtra | Nagpur |
| MS2 | Maharashtra | Nagpur |
| O1 | Odisha | Bhubaneswar |
| O3 | Odisha | Bhubaneswar |
| J1 | Rajasthan | Jaipur |
| J2 | Rajasthan | Jaipur |
| J3 | Rajasthan | Jaipur |
| J4i | Rajasthan | Jaipur |
| J5 | Rajasthan | Jaipur |
| U1 | Rajasthan | Udaipur |
| U2 | Rajasthan | Udaipur |
| TN1 | Tamil Nadu | Chennai |
| TN2 | Tamil Nadu | Chennai |
| TN3 | Tamil Nadu | Chennai |
| JNP1 | Uttar Pradesh | Jaunpur |
| JNP2 | Uttar Pradesh | Jaunpur |
| V2 | Uttar Pradesh | Varanasi |
| V3 | Uttar Pradesh | Varanasi |
| VMS1 | Uttar Pradesh | Varanasi |
| VMS2 | Uttar Pradesh | Varanasi |
| VMS3 | Uttar Pradesh | Varanasi |
| VMS4 | Uttar Pradesh | Varanasi |
| VMS5 | Uttar Pradesh | Varanasi |
| D1 | Uttarakhand | Dehradun |
| D2 | Uttarakhand | Dehradun |

**Table 1.** Frequency of occurrence of different bacterial species (in %)

| **Species** | **Category** | **Frequency** | **Average heterozygosity** | **Number of SNPs** |
| --- | --- | --- | --- | --- |
| *Acinetobacter baumannii* | Non-probiotic | 30% | 0.38 | 10 |
| *Bacillus cereus* | Pathogenic | 15% | 0.56 | 21 |
| *Enterococcus faecium* | Probiotic | 24% | 0 | 1 |
| *Escherichia coli* | Pathogenic | 88% | 0.06 | 9 |
| *Lactobacillus acidophilus* | Probiotic | 18% | 0 | 1 |
| *Lactobacillus delbrueckii subsp bulgaricus* | Probiotic | 100% | 0.07 | 386 |
| *Lactobacillus helveticus* | Probiotic | 100% | 0.33 | 47 |
| *Lactobacillus kefiranofaciens* | Probiotic | 70% | 0.1 | 7 |
| *Lactococcus lactis subsp cremoris* | Probiotic | 55% | 0.16 | 5 |
| *Lactococcus piscium* | Non-probiotic | 48% | 0.25 | 19 |
| *Limosilactobacillus fermentum* | Probiotic | 33% | 0.27 | 3 |
| *Salmonella enterica serovar typhimurium* | Pathogenic | 76% | 0.15 | 14 |
| *Shigella dysenteriae* | Pathogenic | 79% | 0.12 | 11 |
| *Streptococcus salivarius* | Probiotic | 94% | 0.08 | 9 |
| *Streptococcus thermophilus* | Probiotic | 94% | 0.09 | 76 |
| *Enterococcus faecalis* | Non-probiotic | 0% | NA | 0 |
| *Lacticaseibacillus rhamnosus* | Probiotic | 0% | NA | 0 |
| *Leuconostoc lactis* | Probiotic | 0% | NA | 0 |
| *Leuconostoc mesenteroides subsp mesenteroides* | Probiotic | 0% | NA | 0 |
| *Pediococcus pentosaceus* | Probiotic | 0% | NA | 0 |
| *Staphylococcus aureus* | Pathogenic | 0% | NA | 0 |

**Table S2.** Reference genome assemblies used for mapping of the GBS-SNP reads. NCBI AccessionIDs of the assemblies are provided. The size of each genome assembly or scaffold level assembly is provided in base pairs.

| **Species** | **Accession ID** | **Chromosome/Scaffold** | **Size (in bp)** |
| --- | --- | --- | --- |
| *Acinetobacter baumannii* | ASM82879v1 | Chromosome | 4021920 |
| *Bacillus cereus* | ASM2120v1 | Chromosome | 5419036 |
| *Enterococcus faecalis* | ASM636481v1 | Chromosome | 2845446 |
| *Enterococcus faecium* | ASM628035v1 | Chromosome | 2659111 |
| *Enterococcus faecium* | ASM628035v1 | p_unnamned1 | 207651 |
| *Enterococcus faecium* | ASM628035v1 | p_unnamned2 | 63632 |
| *Escherichia coli* | EPEC-E2348_69-V2_ | Chromosome | 4848621 |
| *Lacticaseibacillus rhamnosus* | 44927_D01_ | Chromosome | 2991048 |
| *Lactobacillus acidophilus* | 44927_E01_ | Chromosome | 1982157 |
| *Lactobacillus delbrueckii* subsp*. bulgaricus* | Lactobacillus_bulgaricus_ACA-DC_87_ | Chromosome | 1856003 |
| *Lactobacillus helveticus* | ASM170209v1 | Chromosome | 2058319 |
| *Lactobacillus kefiranofaciens* | IMG-taxon_2597490363_annotated_assembly | Ga0052897_scaffold00001.1 | 236056 |
| *Lactobacillus kefiranofaciens* | IMG-taxon_2597490363_annotated_assembly | Ga0052897_scaffold00002.2 | 187618 |
| *Lactobacillus kefiranofaciens* | IMG-taxon_2597490363_annotated_assembly | Ga0052897_scaffold00003.3 | 118983 |
| *Lactobacillus kefiranofaciens* | IMG-taxon_2597490363_annotated_assembly | Ga0052897_scaffold00004.4 | 113012 |
| *Lactobacillus kefiranofaciens* | IMG-taxon_2597490363_annotated_assembly | Ga0052897_scaffold00005.5 | 91948 |
| *Lactobacillus kefiranofaciens* | IMG-taxon_2597490363_annotated_assembly | Ga0052897_scaffold00006.6 | 77284 |
| *Lactobacillus kefiranofaciens* | IMG-taxon_2597490363_annotated_assembly | Ga0052897_scaffold00007.7 | 75464 |
| *Lactobacillus kefiranofaciens* | IMG-taxon_2597490363_annotated_assembly | Ga0052897_scaffold00008.8 | 71683 |
| *Lactobacillus kefiranofaciens* | IMG-taxon_2597490363_annotated_assembly | Ga0052897_scaffold00009.9 | 69921 |
| *Lactobacillus kefiranofaciens* | IMG-taxon_2597490363_annotated_assembly | Ga0052897_scaffold00010.10 | 67785 |
| *Lactobacillus kefiranofaciens* | IMG-taxon_2597490363_annotated_assembly | Ga0052897_scaffold00011.11 | 55780 |
| *Lactobacillus kefiranofaciens* | IMG-taxon_2597490363_annotated_assembly | Ga0052897_scaffold00012.12 | 55200 |
| *Lactobacillus kefiranofaciens* | IMG-taxon_2597490363_annotated_assembly | Ga0052897_scaffold00013.13 | 49568 |
| *Lactobacillus kefiranofaciens* | IMG-taxon_2597490363_annotated_assembly | Ga0052897_scaffold00014.14 | 49166 |
| *Lactobacillus kefiranofaciens* | IMG-taxon_2597490363_annotated_assembly | Ga0052897_scaffold00015.15 | 48840 |
| *Lactobacillus kefiranofaciens* | IMG-taxon_2597490363_annotated_assembly | Ga0052897_scaffold00016.16 | 48290 |
| *Lactobacillus kefiranofaciens* | IMG-taxon_2597490363_annotated_assembly | Ga0052897_scaffold00017.17 | 46785 |
| *Lactobacillus kefiranofaciens* | IMG-taxon_2597490363_annotated_assembly | Ga0052897_scaffold00018.18 | 42967 |
| *Lactobacillus kefiranofaciens* | IMG-taxon_2597490363_annotated_assembly | Ga0052897_scaffold00019.19 | 42001 |
| *Lactobacillus kefiranofaciens* | IMG-taxon_2597490363_annotated_assembly | Ga0052897_scaffold00020.20 | 41293 |
| *Lactobacillus kefiranofaciens* | IMG-taxon_2597490363_annotated_assembly | Ga0052897_scaffold00021.21 | 38583 |
| *Lactobacillus kefiranofaciens* | IMG-taxon_2597490363_annotated_assembly | Ga0052897_scaffold00022.22 | 38076 |
| *Lactobacillus kefiranofaciens* | IMG-taxon_2597490363_annotated_assembly | Ga0052897_scaffold00023.23 | 27120 |
| *Lactobacillus kefiranofaciens* | IMG-taxon_2597490363_annotated_assembly | Ga0052897_scaffold00024.24 | 25615 |
| *Lactobacillus kefiranofaciens* | IMG-taxon_2597490363_annotated_assembly | Ga0052897_scaffold00025.25 | 25377 |
| *Lactobacillus kefiranofaciens* | IMG-taxon_2597490363_annotated_assembly | Ga0052897_scaffold00026.26 | 25261 |
| *Lactobacillus kefiranofaciens* | IMG-taxon_2597490363_annotated_assembly | Ga0052897_scaffold00027.27 | 24554 |
| *Lactobacillus kefiranofaciens* | IMG-taxon_2597490363_annotated_assembly | Ga0052897_scaffold00028.28 | 22724 |
| *Lactobacillus kefiranofaciens* | IMG-taxon_2597490363_annotated_assembly | Ga0052897_scaffold00029.29 | 19525 |
| *Lactobacillus kefiranofaciens* | IMG-taxon_2597490363_annotated_assembly | Ga0052897_scaffold00030.30 | 18393 |
| *Lactobacillus kefiranofaciens* | IMG-taxon_2597490363_annotated_assembly | Ga0052897_scaffold00031.31 | 17886 |
| *Lactobacillus kefiranofaciens* | IMG-taxon_2597490363_annotated_assembly | Ga0052897_scaffold00032.32 | 17825 |
| *Lactobacillus kefiranofaciens* | IMG-taxon_2597490363_annotated_assembly | Ga0052897_scaffold00033.33 | 17582 |
| *Lactobacillus kefiranofaciens* | IMG-taxon_2597490363_annotated_assembly | Ga0052897_scaffold00034.34 | 16971 |
| *Lactobacillus kefiranofaciens* | IMG-taxon_2597490363_annotated_assembly | Ga0052897_scaffold00035.35 | 15659 |
| *Lactobacillus kefiranofaciens* | IMG-taxon_2597490363_annotated_assembly | Ga0052897_scaffold00036.36 | 15450 |
| *Lactobacillus kefiranofaciens* | IMG-taxon_2597490363_annotated_assembly | Ga0052897_scaffold00037.37 | 14018 |
| *Lactobacillus kefiranofaciens* | IMG-taxon_2597490363_annotated_assembly | Ga0052897_scaffold00038.38 | 13940 |
| *Lactobacillus kefiranofaciens* | IMG-taxon_2597490363_annotated_assembly | Ga0052897_scaffold00039.39 | 13722 |
| *Lactobacillus kefiranofaciens* | IMG-taxon_2597490363_annotated_assembly | Ga0052897_scaffold00040.40 | 13544 |
| *Lactobacillus kefiranofaciens* | IMG-taxon_2597490363_annotated_assembly | Ga0052897_scaffold00041.41 | 13337 |
| *Lactobacillus kefiranofaciens* | IMG-taxon_2597490363_annotated_assembly | Ga0052897_scaffold00042.42 | 12872 |
| *Lactobacillus kefiranofaciens* | IMG-taxon_2597490363_annotated_assembly | Ga0052897_scaffold00043.43 | 12243 |
| *Lactobacillus kefiranofaciens* | IMG-taxon_2597490363_annotated_assembly | Ga0052897_scaffold00044.44 | 12095 |
| *Lactobacillus kefiranofaciens* | IMG-taxon_2597490363_annotated_assembly | Ga0052897_scaffold00045.45 | 11986 |
| *Lactobacillus kefiranofaciens* | IMG-taxon_2597490363_annotated_assembly | Ga0052897_scaffold00046.46 | 11740 |
| *Lactobacillus kefiranofaciens* | IMG-taxon_2597490363_annotated_assembly | Ga0052897_scaffold00047.47 | 11658 |
| *Lactobacillus kefiranofaciens* | IMG-taxon_2597490363_annotated_assembly | Ga0052897_scaffold00048.48 | 11563 |
| *Lactobacillus kefiranofaciens* | IMG-taxon_2597490363_annotated_assembly | Ga0052897_scaffold00049.49 | 10929 |
| *Lactobacillus kefiranofaciens* | IMG-taxon_2597490363_annotated_assembly | Ga0052897_scaffold00050.50 | 10747 |
| *Lactobacillus kefiranofaciens* | IMG-taxon_2597490363_annotated_assembly | Ga0052897_scaffold00051.51 | 10562 |
| *Lactobacillus kefiranofaciens* | IMG-taxon_2597490363_annotated_assembly | Ga0052897_scaffold00052.52 | 9361 |
| *Lactobacillus kefiranofaciens* | IMG-taxon_2597490363_annotated_assembly | Ga0052897_scaffold00053.53 | 9031 |
| *Lactobacillus kefiranofaciens* | IMG-taxon_2597490363_annotated_assembly | Ga0052897_scaffold00054.54 | 8204 |
| *Lactobacillus kefiranofaciens* | IMG-taxon_2597490363_annotated_assembly | Ga0052897_scaffold00055.55 | 8146 |
| *Lactobacillus kefiranofaciens* | IMG-taxon_2597490363_annotated_assembly | Ga0052897_scaffold00056.56 | 7488 |
| *Lactobacillus kefiranofaciens* | IMG-taxon_2597490363_annotated_assembly | Ga0052897_scaffold00057.57 | 7312 |
| *Lactobacillus kefiranofaciens* | IMG-taxon_2597490363_annotated_assembly | Ga0052897_scaffold00058.58 | 6472 |
| *Lactobacillus kefiranofaciens* | IMG-taxon_2597490363_annotated_assembly | Ga0052897_scaffold00059.59 | 6413 |
| *Lactobacillus kefiranofaciens* | IMG-taxon_2597490363_annotated_assembly | Ga0052897_scaffold00060.60 | 5874 |
| *Lactobacillus kefiranofaciens* | IMG-taxon_2597490363_annotated_assembly | Ga0052897_scaffold00061.61 | 5316 |
| *Lactobacillus kefiranofaciens* | IMG-taxon_2597490363_annotated_assembly | Ga0052897_scaffold00062.62 | 5041 |
| *Lactobacillus kefiranofaciens* | IMG-taxon_2597490363_annotated_assembly | Ga0052897_scaffold00063.63 | 5028 |
| *Lactobacillus kefiranofaciens* | IMG-taxon_2597490363_annotated_assembly | Ga0052897_scaffold00064.64 | 4915 |
| *Lactobacillus kefiranofaciens* | IMG-taxon_2597490363_annotated_assembly | Ga0052897_scaffold00065.65 | 4867 |
| *Lactobacillus kefiranofaciens* | IMG-taxon_2597490363_annotated_assembly | Ga0052897_scaffold00066.66 | 4676 |
| *Lactobacillus kefiranofaciens* | IMG-taxon_2597490363_annotated_assembly | Ga0052897_scaffold00067.67 | 4644 |
| *Lactobacillus kefiranofaciens* | IMG-taxon_2597490363_annotated_assembly | Ga0052897_scaffold00068.68 | 4347 |
| *Lactobacillus kefiranofaciens* | IMG-taxon_2597490363_annotated_assembly | Ga0052897_scaffold00069.69 | 3198 |
| *Lactobacillus kefiranofaciens* | IMG-taxon_2597490363_annotated_assembly | Ga0052897_scaffold00070.70 | 3195 |
| *Lactobacillus kefiranofaciens* | IMG-taxon_2597490363_annotated_assembly | Ga0052897_scaffold00072.72 | 2576 |
| *Lactobacillus kefiranofaciens* | IMG-taxon_2597490363_annotated_assembly | Ga0052897_scaffold00071.71 | 2563 |
| *Lactobacillus kefiranofaciens* | IMG-taxon_2597490363_annotated_assembly | Ga0052897_scaffold00073.73 | 2473 |
| *Lactobacillus kefiranofaciens* | IMG-taxon_2597490363_annotated_assembly | Ga0052897_scaffold00074.74 | 2375 |
| *Lactobacillus kefiranofaciens* | IMG-taxon_2597490363_annotated_assembly | Ga0052897_scaffold00075.75 | 2307 |
| *Lactobacillus kefiranofaciens* | IMG-taxon_2597490363_annotated_assembly | Ga0052897_scaffold00076.76 | 2290 |
| *Lactobacillus kefiranofaciens* | IMG-taxon_2597490363_annotated_assembly | Ga0052897_scaffold00077.77 | 2198 |
| *Lactobacillus kefiranofaciens* | IMG-taxon_2597490363_annotated_assembly | Ga0052897_scaffold00078.78 | 2068 |
| *Lactobacillus kefiranofaciens* | IMG-taxon_2597490363_annotated_assembly | Ga0052897_scaffold00079.79 | 1933 |
| *Lactobacillus kefiranofaciens* | IMG-taxon_2597490363_annotated_assembly | Ga0052897_scaffold00080.80 | 1547 |
| *Lactobacillus kefiranofaciens* | IMG-taxon_2597490363_annotated_assembly | Ga0052897_scaffold00081.81 | 1312 |
| *Lactobacillus kefiranofaciens* | IMG-taxon_2597490363_annotated_assembly | Ga0052897_scaffold00082.82 | 1270 |
| *Lactobacillus kefiranofaciens* | IMG-taxon_2597490363_annotated_assembly | Ga0052897_scaffold00083.83 | 1249 |
| *Lactobacillus kefiranofaciens* | IMG-taxon_2597490363_annotated_assembly | Ga0052897_scaffold00084.84 | 1062 |
| *Lactococcus lactis* subsp. *cremoris* | ASM46895v1_ | Chromosome | 2427048 |
| *Lactococcus piscium* | L_piscium_ | I | 2394138 |
| *Lactococcus piscium* | L_piscium_ | II | 55671 |
| *Lactococcus piscium* | L_piscium_ | III | 53257 |
| *Leuconostoc lactis* | ASM169814v1_ | Chromosome | 1737502 |
| *Leuconostoc lactis* | ASM169814v1_ | pWK40-1 | 20388 |
| *Leuconostoc lactis* | ASM169814v1_ | pWK40-2 | 19726 |
| *Leuconostoc lactis* | ASM169814v1_ | pWK40-3 | 10453 |
| *Lactococcus lactis* subsp*. cremoris* | ASM214823v1_ | Chromosome | 1891719 |
| *Leuconostoc mesenteroides* subsp*. mesenteroides* | ASM214823v1_ | pLm1 | 31104 |
| *Leuconostoc mesenteroides* subsp. *mesenteroides* | ASM214823v1_ | pLm2 | 15285 |
| *Leuconostoc mesenteroides* subsp*. mesenteroides* | ASM214823v1_ | pLm3 | 8768 |
| *Limosilactobacillus fermentum* | Lf_IMDO_130101_ | Chromosome | 2089202 |
| *Pediococcus pentosaceus* | ASM1450v1_ | Chromosome | 1832387 |
| *Salmonella enterica* subsp. *enterica* serovar Typhimurium | ASM635180v1 | Chromosome | 4813658 |
| *Salmonella enterica* subsp. *enterica* serovar Typhimurium | ASM635180v1 | pCFSAN074387 | 93930 |
| *Salmonella enterica* subsp. *enterica* serovar Typhimurium | ASM1200v1 | pSD197_spA | 8953 |
| *Salmonella enterica* subsp. *enterica* serovar Typhimurium | ASM1200v1 | pSD1_197 | 182726 |
| *Shigella dysenteriae* | ASM1200v1 | Chromosome | 4369232 |
| *Staphylococcus aureus* | 40961_D01_ | Chromosome | 2563627 |
| *Streptococcus salivarius* | 42197_C01_ | Chromosome | 2206150 |
| *Streptococcus thermophilus* | TH1477VE_ | chr | 1834035 |
| *Streptococcus thermophilus* | TH1477VE_ | scaffold48 | 18808 |
| *Streptococcus thermophilus* | TH1477VE_ | scaffold44 | 5601 |
| *Streptococcus thermophilus* | TH1477VE_ | scaffold46 | 4943 |
| *Streptococcus thermophilus* | TH1477VE_ | scaffold55 | 4283 |
| *Streptococcus thermophilus* | TH1477VE_ | scaffold45 | 4070 |
| *Streptococcus thermophilus* | TH1477VE_ | scaffold54 | 3889 |
| *Streptococcus thermophilus* | TH1477VE_ | scaffold50 | 3827 |
| *Streptococcus thermophilus* | TH1477VE_ | scaffold51 | 2535 |
| *Streptococcus thermophilus* | TH1477VE_ | scaffold42 | 1729 |
| *Streptococcus thermophilus* | TH1477VE_ | scaffold53 | 1504 |
| *Streptococcus thermophilus* | TH1477VE_ | scaffold56 | 1069 |
| *Streptococcus thermophilus* | TH1477VE_ | scaffold43 | 1033 |
| *Streptococcus thermophilus* | TH1477VE_ | scaffold52 | 933 |
| *Streptococcus thermophilus* | TH1477VE_ | scaffold49 | 684 |
| *Streptococcus thermophilus* | TH1477VE_ | scaffold47 | 613 |

**Table S3.** Occurrence of different bacterial species in dahi samples. “1” denotes presence of a species and “0” denotes absence of the species. The last column shows total number of species present in the sample. The last row shows the total number of dahi samples in which the species was present.

| **Species** | *Acinetobacter baumannii* | *Bacillus cereus* | *Enterococcus faecium* | *Escherichia coli* | *Lactobacillus acidophilus* | *Lactobacillus delbrueckii* subsp *bulgaricus* | *Lactobacillus helveticus* | *Lactobacillus kefiranofaciens* | *Lactococcus lactis* subsp *cremoris* | *Lactococcus piscium* | *Limosilactobacillus fermentum* | *Salmonella enterica serovar typhimurium* | *Shigella dysenteriae* | *Streptococcus salivarius* | *Streptococcus thermophilus* | **Total count** |
| --- | --- | --- | --- | --- | --- | --- | --- | --- | --- | --- | --- | --- | --- | --- | --- | --- |
| **D1** | 0 | 0 | 1 | 1 | 1 | 1 | 1 | 1 | 1 | 0 | 0 | 1 | 1 | 1 | 1 | **11** |
| **D2** | 0 | 0 | 1 | 0 | 0 | 1 | 1 | 1 | 0 | 1 | 0 | 0 | 0 | 1 | 1 | **7** |
| **J1** | 0 | 0 | 0 | 1 | 0 | 1 | 1 | 0 | 0 | 0 | 0 | 0 | 1 | 1 | 1 | **6** |
| **J2** | 0 | 0 | 0 | 1 | 0 | 1 | 1 | 1 | 1 | 1 | 0 | 0 | 1 | 1 | 1 | **9** |
| **J3** | 0 | 0 | 0 | 1 | 0 | 1 | 1 | 1 | 1 | 1 | 0 | 0 | 0 | 1 | 1 | **8** |
| **J4 i** | 0 | 0 | 1 | 1 | 0 | 1 | 1 | 1 | 1 | 1 | 0 | 1 | 0 | 1 | 1 | **10** |
| **J5** | 0 | 0 | 0 | 1 | 0 | 1 | 1 | 1 | 0 | 1 | 0 | 1 | 1 | 1 | 1 | **9** |
| **JNP1** | 1 | 0 | 1 | 1 | 1 | 1 | 1 | 1 | 0 | 0 | 0 | 1 | 1 | 1 | 1 | **11** |
| **JNP2** | 0 | 0 | 0 | 1 | 0 | 1 | 1 | 0 | 0 | 0 | 1 | 1 | 1 | 1 | 1 | **8** |
| **K1** | 0 | 0 | 0 | 1 | 1 | 1 | 1 | 1 | 0 | 0 | 1 | 1 | 1 | 0 | 0 | **8** |
| **K4** | 0 | 0 | 0 | 1 | 1 | 1 | 1 | 1 | 0 | 0 | 1 | 1 | 1 | 0 | 0 | **8** |
| **L** | 1 | 0 | 0 | 0 | 0 | 1 | 1 | 0 | 1 | 1 | 1 | 1 | 1 | 1 | 1 | **10** |
| **MS** | 0 | 0 | 0 | 1 | 0 | 1 | 1 | 1 | 0 | 1 | 1 | 0 | 1 | 1 | 1 | **9** |
| **MS2** | 0 | 0 | 0 | 1 | 0 | 1 | 1 | 1 | 0 | 0 | 1 | 0 | 0 | 1 | 1 | **7** |
| **O1** | 0 | 0 | 0 | 1 | 0 | 1 | 1 | 0 | 1 | 1 | 1 | 1 | 1 | 1 | 1 | **10** |
| **O3** | 0 | 0 | 0 | 1 | 0 | 1 | 1 | 1 | 1 | 0 | 0 | 1 | 0 | 1 | 1 | **8** |
| **P3** | 1 | 0 | 1 | 1 | 0 | 1 | 1 | 0 | 0 | 0 | 0 | 1 | 1 | 1 | 1 | **9** |
| **P4** | 1 | 0 | 1 | 1 | 0 | 1 | 1 | 0 | 1 | 1 | 0 | 1 | 1 | 1 | 1 | **11** |
| **P5** | 1 | 0 | 0 | 1 | 0 | 1 | 1 | 0 | 1 | 0 | 0 | 1 | 1 | 1 | 1 | **9** |
| **P6** | 1 | 1 | 0 | 1 | 0 | 1 | 1 | 0 | 0 | 1 | 0 | 1 | 1 | 1 | 1 | **10** |
| **P7** | 1 | 1 | 0 | 1 | 0 | 1 | 1 | 0 | 1 | 1 | 0 | 1 | 1 | 1 | 1 | **11** |
| **TN1** | 1 | 1 | 1 | 1 | 0 | 1 | 1 | 1 | 1 | 1 | 0 | 1 | 1 | 1 | 1 | **13** |
| **TN2** | 0 | 1 | 0 | 1 | 0 | 1 | 1 | 1 | 1 | 1 | 0 | 0 | 1 | 1 | 1 | **10** |
| **TN3** | 0 | 1 | 0 | 1 | 0 | 1 | 1 | 1 | 1 | 1 | 1 | 1 | 1 | 1 | 1 | **12** |
| **U1** | 0 | 0 | 0 | 1 | 1 | 1 | 1 | 1 | 0 | 0 | 1 | 1 | 1 | 1 | 1 | **10** |
| **U2** | 0 | 0 | 0 | 1 | 1 | 1 | 1 | 1 | 1 | 1 | 1 | 1 | 1 | 1 | 1 | **12** |
| **V2** | 1 | 0 | 0 | 1 | 0 | 1 | 1 | 1 | 0 | 0 | 0 | 1 | 1 | 1 | 1 | **9** |
| **V3** | 0 | 0 | 0 | 1 | 0 | 1 | 1 | 1 | 1 | 1 | 1 | 1 | 1 | 1 | 1 | **11** |
| **VMS1** | 0 | 0 | 1 | 1 | 0 | 1 | 1 | 1 | 1 | 0 | 0 | 1 | 1 | 1 | 1 | **10** |
| **VMS2** | 1 | 0 | 0 | 1 | 0 | 1 | 1 | 1 | 1 | 0 | 0 | 1 | 1 | 1 | 1 | **10** |
| **VMS3** | 0 | 0 | 0 | 1 | 0 | 1 | 1 | 1 | 1 | 0 | 0 | 1 | 1 | 1 | 1 | **9** |
| **VMS4** | 0 | 0 | 0 | 0 | 0 | 1 | 1 | 1 | 0 | 0 | 0 | 1 | 0 | 1 | 1 | **6** |
| **VMS5** | 0 | 0 | 0 | 0 | 0 | 1 | 1 | 0 | 0 | 0 | 0 | 0 | 0 | 1 | 1 | **4** |
| **Total** | **10** | **5** | **8** | **29** | **6** | **33** | **33** | **23** | **18** | **16** | **11** | **25** | **26** | **31** | **31** |  |

**Figure S1**: UPGMA clustering of curd samples based on presence/ absence of different bacterial species

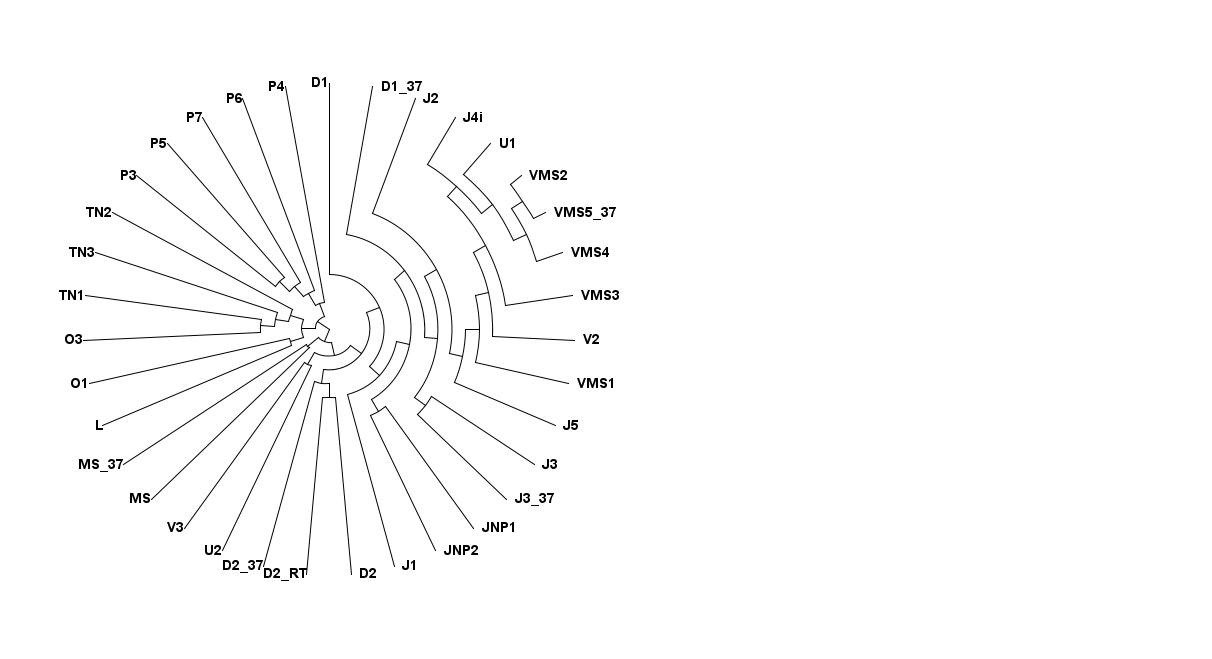

**Figure S2**: Genetic relatedness of *Streptococcus thermophilus* strains. A total of 207 GBS-SNPs were used to generate the UPGMA dendrogram based on identity-by-state.

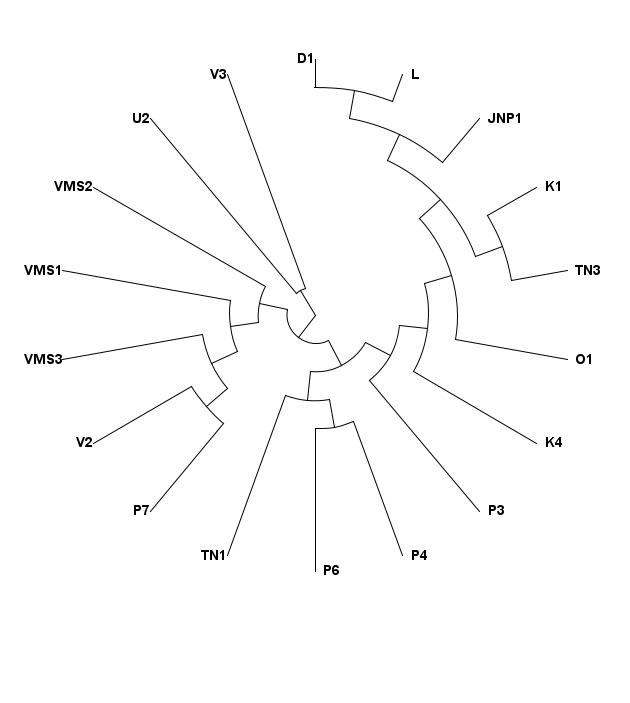

**Figure S3**: Genetic relatedness of *Salmonella enterica* serovar *typhimurium* strains. A total of 50 GBS-SNPs were used to generate the UPGMA dendrogram based on identity-by-state.
